# Evaluating *de novo* assembly and binning strategies for time-series drinking water metagenomes

**DOI:** 10.1101/2021.07.11.451960

**Authors:** Solize Vosloo, Linxuan Huo, Christopher L. Anderson, Zihan Dai, Maria Sevillano, Ameet Pinto

**Author notes:** Corresponding author: Ameet Pinto.

## Abstract

Reconstructing microbial genomes from metagenomic short-read data can be challenging due to the unknown and uneven complexity of microbial communities. This complexity encompasses highly diverse populations which often includes strain variants. Reconstructing high-quality genomes is a crucial part of the metagenomic workflow as subsequent ecological and metabolic inferences depend on their accuracy, quality, and completeness. In contrast to microbial communities in other ecosystems, there has been no systematic assessment of genome-centric metagenomic workflows for drinking water microbiomes. In this study, we assessed the performance of a combination of assembly and binning strategies for time-series drinking water metagenomes that were collected over a period of 6 months. The goal of this study was to identify the combination of assembly and binning approaches that results in high quality and quantity metagenome-assembled genomes (MAGs), representing most of the sequenced metagenome. Our findings suggest that the metaSPAdes co-assembly strategies had the best performance as they resulted in larger and less fragmented assemblies with at least 85% of the sequence data mapping to contigs greater than 1kbp. Furthermore, a combination of metaSPAdes co-assembly strategies and MetaBAT2 produced the highest number of medium-quality MAGs while capturing at least 70% of the metagenomes based on read recruitment. Utilizing different assembly/binning approaches also assist in the reconstruction of unique MAGs from closely related species that would have otherwise collapsed into a single MAG using a single workflow. Overall, our study suggests that leveraging multiple binning approaches with different metaSPAdes co-assembly strategies may be required to maximize the recovery of good-quality MAGs, which more accurately capture the microbial diversity of drinking water samples.

## Introduction

Advances in high-throughput sequencing technologies have enabled characterization of microbial communities without the need for cultivation (1). This has greatly facilitated our understanding of microbial communities that inhabit a range of natural and engineered ecosystems. Two high-throughput sequencing technologies commonly used to characterize microbial communities includes gene-targeted assays that uses universal genes/regions (i.e., 16S rRNA, 18S rRNA and internal transcribe spacer region for bacteria/archaea, eukaryotes, and fungi, respectively) and short read shotgun DNA sequencing (i.e., metagenomics) (1–4). Other emerging sequencing approaches includes synthetic- and single-molecule long-read sequencing for both gene-targeted and metagenomic assays (5, 6). Gene-targeted assays provide valuable insights into the compositional and structural profiles of microbial communities in a fast and cost-effective manner; however, this approach is limited by challenges related to primer selection and amplification bias (7). Furthermore, taxonomic classification in gene-targeted assays is based on a fragment of a singular conserved universal marker gene that permits little resolution beyond the genus level and does not allow for the direct analysis of a microbial community’s metabolic capabilities (8). In some instances, putative functional assignment is possible when using gene-targeted assays; however, this requires the availability of a curated taxonomic databases and classification beyond the genus level (9). Limitations of gene-targeted assays can be overcome by utilizing genome-resolved metagenomics (10, 11). Genome-resolved metagenomics encompasses *de novo* assembly of short high-throughput paired-end reads into longer contiguous sequences (contigs) and subsequent reconstruction of metagenome-assembled genomes (MAGs) through clustering (or binning) of contigs based on nucleotide composition and differential coverage (12, 13). This approach offers improved taxonomic and functional potential analysis, as well as the characterization of novel microorganisms using phylogenetic analysis (14).

De novo assembly and reconstruction of MAGs from short-read metagenomic data can be challenging due to sequencing errors, repeats, depth of sequencing coverage, and the presence of strain variants (15, 16). These challenges influence the performance of assemblers as it creates unresolved ambiguities in the reconstructed contigs, leading to erroneous and/or fragmented assemblies. Reconstructing high-quality MAGs is a crucial part of the genome-centric metagenomic workflow as subsequent taxonomic, metabolic, and ecological inferences depend on the accuracy, quality, and completeness of genomes. Studies have attempted to optimize the recovery of high-quality assemblies and MAGs by benchmarking metagenomic software for assembly, binning, and taxonomic classification (16–19). However, owing to the unknown complexity of varying environmental sample types, systematic evaluation of metagenomic workflows is required as tool selection depends on the complexity of the biological sample and the availability of computational recourses (18).

Genomes are often reconstructed by assembling all the samples together (co-assembly) or creating individual assemblies. Co-assembly is a computationally intensive approach that involves the pooling of multiple metagenomes, which allow for greater sequence depth and coverage as well as leveraging differential coverage of microorganisms across genomes for genome binning. While this assembly approach can facilitate the identification of populations that are present at lower abundances, it can also result in ambiguous and/or fragmented assemblies when strain level variability is high (20, 21). In contrast, single sample assemblies are computationally less intensive and are often used to reconstruct genomes of larger datasets and to preserve strain variation between different samples (22). It has also been shown that single sample assembly produce more non-redundant high-quality MAGs and enables the reconstructions of genomes with similar phylogenetic placement compared to co-assembled genomes (14). However, lower sequence depth and thus lower coverage resulting from single-sample assembly in addition to the lack of differential coverage information, makes genome reconstructions difficult when using this assembly approach as coverage heuristics that are used to accurately disentangle repetitive sequences and differentiate between strain variants cannot be properly applied.

Advances in our understanding of the drinking water microbiome have been greatly facilitated by the application of genome-resolve metagenomics (23–26). Despite this progress, inferences of microbial community dynamics in drinking water systems (DWS) have been restrained by the limited availability of longitudinal metagenomic datasets as most previous work was done utalizing gene-targeted assays in studies that were short (i.e., few time points), gapped (i.e., missing time points) and/or implemented over multiple spatio-temporal scales (27–29). Longitudinal datasets are preferred over cross-sectional studies (1) as they offer unique insights into the stability and dynamics of microbial communities. This is because information leveraged from these datasets can reveal periodic patterns that can be used in predictive modeling, describe irregularities in response to abrupt environmental perturbations, and capture temporal variation of microbial interactions (30). Currently, there is little work on how to best leverage the unique properties of time-series metagenomic data for DWS. Thus, the overall objective of this study was to evaluate the performance of a combination of *de novo* assembly and binning algorithms for time-series metagenomic data for drinking water microbial communities. Our goal was to identify an ideal combination of assembly and binning strategies that can allow for high quality metagenomic assemblies and MAGs that maximally capture the sequenced metagenomes.

## Materials and Methods

### Sample Collection

Samples (*n* = 12) were collected over a period of 6 months from a tap in a commercial building located in Boston, MA (United States) (Table S1). Prior to sample collection, the system was flushed for at least 30 min at a flow rate ranging between 3.0 and 3.3 l.min^-1^ and then approximately 1,500 ml of tap water was collected for microbial community analysis in a sterile (by autoclaving) 2 L DURAN® GLS 80® wide mouth borosilicated glass bottle (DURAN®, Cat. No.: 1112715). An additional 500 ml sample was collected in parallel in sterile 2 × 250 ml DURAN® GLS 80® wide mouth borosilicated glass bottles (DURAN®, Cat. No.: 218603656) for chemical analysis. Samples for microbial community analysis were filtered immediately through Sterivex-GP Pressure Filter Units (EMD Millipore, Cat. No.: SVGP01050) containing a 0.22μm polyethersulfone (PES) filter membrane, using the Geotech Geopump™ Series II peristaltic pump (Geotech Environmental Equipment, Inc., Cat. No.: 91350113) and sterile SZ 15 Geotech silicone tubing (Geotech Environmental Equipment, Inc., Cat. No.: 77050000). Following filtration, the exterior of the filter unit was cleaned with an 70% ethanol (Fisher Scientific, Cat. No.: A962F) soaked Kimwipe (Kimberly-Clark Professional™, Cat. No.: 34120) and then transferred to a 50 ml Falcon tube (Corning, Cat. No.: 362070) and stored at -80°C until further analysis.

### Water chemistry characterization

Water quality parameters (i.e., temperature, pH, conductivity, and dissolved oxygen) were measured using the Orion Star™ A325 pH/Conductivity Portable Multiparameter Meter (Thermo Scientific™, Cat. No.: STARA3250). Total chlorine was measured using USEPA approved HACH Method 8167 with DPD Total Chlorine Reagent Powder Pillows (HACH, Cat. No.: 2105669). Reactive orthophosphate was measured using USEPA approved HACH Method 8048 with PhosVer®3 Phosphate Reagent Powder Pillows (HACH, Cat. No.: 2106028). Nitrogen species, including ammonium, nitrate and nitrate were measured using the Nitrogen-Ammonia Reagent Set (Method 10023, HACH, Cat. No.: 2604545), NitraVer X Nitrogen-Nitrate Reagent Set (Method 10020 HACH, Cat. No.: 2605345), and NitriVer 3 TNT Reagent Set (Method 10019, Cat. No.: 2608345), respectively. All HACH measurements were performed in triplicate on the DR1900 Portable Spectrophotometer (HACH, Cat. No.: DR190001H) (Table S1).

### Flow cytometric analysis

Standard flow cytometric measurements (FCM) were performed as described previously (31, 32). Briefly, samples were quenched with 10mM sodium thiosulfate (1% (v/v)) (Alfa Aesar™, Cat. No.: AA35645K2) and then pre-heated at 37°C for 3 min, stained with SYBR Green I (SG) (Invitrogen™, Cat. No.: S7585) (1:100 diluted in 10 mM Tris-HCl (pH 8.5, Bioworld, Cat. No: NC1213695)) at 10 μl.ml^-1^ or SG combined with propidium iodide (PI) (Molecular Probes™, Cat. No.: P3566) (3uM final concentration) at 12 μl.ml^-1^ and incubated in the dark at 37°C for 10 min. Five negative controls consisting of (i) unstained UltraPure™ DNase/RNase-Free Distilled Water (Thermo Fisher Scientific, Cat. No.: 10977015), (ii) SG stained UltraPure™ DNase/RNase-Free Distilled Water, (iii) SGPI stained UltraPure™ DNase/RNase-Free Distilled Water, (iv) SG stained 0.22μm filtered tap water sample, and (v) SGPI stained 0.22μm filtered tap water sample were processed identically and in parallel with the samples. FCM were performed on 50 μl sample in triplicate at a pre-set flow rate of 66 μl.min^-1^ using a BD Accuri® C6 flow cytometer (BD Accuri® cytometers, Belgium) which is equipped with a 50mW solid state laser emitting light at a fixed wavelength of 488 nm. Green and red fluorescent intensity was collected at FL1 = 533 ± 30 nm and FL3 > 670 nm, respectively, along with sideward and forward scatter light intensities. Data were processed with the BD Accuri CFlow® software that permits electronic gating to separate the positive signals from instrumental and sample background noise on a two-parameter density plot (33). A trigger/threshold of 1,000 was applied on the green fluorescence channel (FL1). No compensation was used.

### Sample Processing and DNA Extraction

Prior to extraction, the bead constituents (i.e., ceramic and silica spheres, and glass bed) contained within the 2 ml Lysing Matrix E tubes (MP Biomedicals, Cat. No.: 116914100) were aseptically transferred into sterile 1.5 ml microcentrifuge tubes (Eppendorf, Cat. No.: 022431021) (34). Removal of these components was necessary to ensure that the processed PES filter membranes from the Sterivex-GP Pressure Filter Units are fully immersed in solution during the enzymatic and chemical treatment steps of the DNA extraction protocol. The PES filter membrane with harvested microbial biomass was aseptically removed from the Sterivex-GP Pressure Filter Unit and cut into smaller pieces on the surface of a petri dish (Fisher Scientific, Cat. No.: FB0875712) using a sterile scalpel (Fisher Scientific, Cat. No.: 08-920B) and then transferred into the emptied 2 ml Lysing Matrix E tubes using a sterile tweezer (Fisher Scientific, Cat. No: 22327379). DNA extractions were performed using a modified version of the DNeasy PowerWater Kit® (QIAGEN, Cat. No.: 14900-50-NF or 14900-100-NF) protocol that utilizes enzymatic, chemical, and mechanical lysis strategies to enhance recovery of DNA from drinking water samples (34). Briefly, filter cuttings contained in the 2 ml Lysing Matrix E tubes were submerged in 294 µl 10X Tris-EDTA (100 mM Tris, 10 mM EDTA, pH 8.0, G-Biosciences, Cat. No: 501035446) and 6 μl lysozyme solution (50 mg.ml^-1^, Thermo Fisher Scientific, Cat. No.: 90082) and incubated for 60 min at 37°C with light mixing at 300 rpm using the Eppendorf ThermoMixer® C (Eppendorf, Cat. No.: 2231000680). Subsequently, the tubes were supplemented with 300 μl pre-warmed (55°C) PW1 solution, provided with the DNeasy PowerWater Kit®, and 30 μl Proteinase K (20 mg.ml^-1^, Thermo Fisher Scientific, Cat. No.: AM2546), vortexed and incubated for 30 min at 56°C with light mixing at 300 rpm using the Eppendorf ThermoMixer® C. After incubation, the bead constituents initially transferred to the sterile 1.5 ml microcentrifuge tubes were aseptically transferred back to the Lysing Matrix E tubes. The tubes were then supplemented with 630 μl chloroform/isoamyl alcohol (24:1, pH 8, Acros Organics, Cat. No.: 327155000) and bead beat at setting 6 for 40 sec using the FastPrep-24™ Classic Instrument (MP Biomedicals, Cat. No.: 116004500). The resulting homogenized mixture was then subjected to centrifugation at 14 000 x g for 10 min at 4°C using the Eppendorf® Centrifuge 5424R (Cat. No.: 5404000332). After centrifugation, the aqueous phase (600 - 650 μl) was transferred to a sterile 1.5 ml microcentrifuge tube. Exactly 600 μl of the aqueous phase was used as starting material on the QIACube System (QIAGEN, Cat. No.: 9001882) to purify DNA according to the manufacture instructions using the DNeasy PowerWater Kit® protocol. Three negative controls consisting of a reagent blank (C01) and two filter blanks (i.e., unused PES membrane filters (C02) and PES membrane filters treated with autoclave deionized water (C03)) were processed identically and in parallel with the samples. The extracted DNA was quantified in duplicate using the Qubit™ dsDNA High Sensitivity (HS) Assay Kit (Thermo Fisher Scientific, Cat. No.: Q32851) with the Qubit™ 4 Fluorometer (Thermo Fisher Scientific, Cat. No.: Q33238) (Table S2). All DNA extracts (50 μl) were stored at -80°C until further analysis.

### Quantitative PCR

The quantitative PCR (qPCR) assay was performed on a QuantStudio™ 3 Real-Time PCR System (ThermoFisher Scientific Cat. no. A28567) in a 20 μl reaction mixture consisting of Luna® Universal qPCR Master Mix (New England Biolabs, Inc., Cat. No.: NC1276266), forward and reverse primer pairs (F515-GTGCCAGCMGCCGCGGTAA and R806-GGACTACHVGGGTWTCTAAT, respectively) (35), UltraPure™ DNase/RNase-Free Distilled Water (Thermo Fisher Scientific, Cat. No.: 10977015) and 1:10 diluted DNA template. Reactions were prepared in triplicate in a 96-well optical plate using the epMOTION® M5073 automated liquid handling system (Eppendorf, Cat. no. 5073000205D). qPCR conditions were as follow: 1 min at 95°C, and then 40 cycles consisting of 15 sec at 95°C, 15 sec at 50°C and 1 min at 72°C. A calibration curve with standards ranging from 10^2^ - 10^8^ copies of 16S rRNA gene of *Nitrosomonas europaea* for total bacteria assay were generated. The calibration curve for 16S rRNA copies was linear (R^2^ = 0.997) over 7 orders of magnitude with a high PCR efficiency (100%).

### Metagenomic Sequencing

Sequencing libraries were prepared using the Ovation® Ultralow DNA-Seq Library Preparation Kit (NuGEN, Cat. No.: 0344NB). Metagenomic sequencing was performed on one SP lane of the NovaSeq 6000 sequencing system (Illumina) at the Roy J. Carver Biotechnology Centre at the University of Illinois Urbana-Champaign (UIUC) Sequencing Core (Champaign, IL, United States).

### Sequence Processing

#### Pre-processing

Processing of sequencing data was done using the workflow outlined in Figure S1. Initial quality control of FASTQ files were performed using fastp v0.20.0 (36) with parameters: --trim_poly_x, --qualified_quality_phred 20, --length_required 20. The UniVec_Core database from NCBI (ftp://ftp.ncbi.nih.gov/pub/UniVec/) was subsequently used to screen for contaminant sequences (e.g., phix sequencing control used as sequencing control and sequencing adapters) by mapping the reads from each sample against the UniVec_Core database using BWA-MEM v0.7.17 (37) and then filtering reads in proper pair and supplementary alignments using samtools v1.9 (38) with parameters: -hbS -F2 -F2048. BAM files were subsequently sorted using the sort function of samtools v1.9 and then quality filtered forward and reverse FASTQ files were extracted from sequence alignments in sorted BAM format using the bamtofastq function of bedtools v2.29.2 (https://bedtools.readthedocs.io/en/latest/). The quality filtered FASTQ files were analyzed using Nonpareil v3.303 (39) in kmer mode to estimate the coverage and to predict the number of sequences required to achieve “near complete” coverage. Nonpareil curves were generated in R (40) using the function Nonpareil.set of Nonpareil v3.3.4.

#### MASH distance and k-means clustering

MASH v2.2.2 (41) was used to estimate read-based dissimilarity between samples using the quality filtered FASTQ files. For this, forward and reverse quality filtered FASTQ files of each sample were interleaved using interleafq v1.0 (https://github.com/quadram-institutebioscience/interleafq) and then the sketch function was used to convert the interleaved quality filtered FASTQ files of each sample into a MinHash sketch with parameters: s = 100,000 and k = 21. The dist function was subsequently used for pairwise comparisons between samples based on Jaccard indices; thereby comparing the fraction shared k-mers between samples. K-means clustering on MASH distances was performed to partition samples into clusters with the nearest mean (Figure S2). For this, the MASH-distance matrix was imported into R and the function fviz_nbclust of factoextra v1.0.7 (https://www.rdocumentation.org/packages/factoextra) was used to determine and visualize the optimal number of clusters (or k groups) using the average silhouette method with 999 Monte Carlo iterations. The MASH-distance matrix was subsequently clustered by the k-means method using kmeans of the stats package v3.6.2 (https://www.rdocumentation.org/packages/stats/versions/3.6.2/topics/kmeans) and then visualized using fviz_cluster of factoextra v1.0.7.

#### Metagenomic assembly and binning

The performance of a combination of assembly (metaSPAdes v.3.13.1(42) and MEGAHIT v.1.2.9 (43)), binning (CONCOCT v.1.1.1 (13), MetaBAT v.2.12.1 (12), MaxBin v.2.2.4 (44)) and bin aggregating software (DAS Tool v.1.1.0 (45)) were evaluated using four assembly strategies, including individual assembly and three co-assembly approaches, i.e., co-assembly with all samples, MASH distance-based assembly, and time-discrete assembly. This resulted in 32 combinations of assembler, assembly strategy, and binning approaches (Table S2). For MASH distance-based assemblies, three co-assemblies consisting of pooled samples that were identified using pair-wise MASH dissimilarity indices and kmeans clustering were identified: (i) BW003 + BW015 + BW030 + BW060, (ii) BW075 + BW090 + BW105, and (iii) BW120 + BW135 + BW150 + BW165 (Table S2). Samples pooled and co-assembled for time-discrete assembly consisted of eleven combinations representing paired samples of successive sampling points: (i) BW003 + BW015, (ii) BW015 + BW030, (iii) BW030 + BW045, (iv) BW045 + BW060, (v) BW060 + BW075, (vi) BW075 + BW090, (vii) BW090 + BW105, (viii) BW105 + BW120, (ix) BW120 + BW135, (x) BW135 + BW150, and (xi) BW150 + BW165. Control samples (i.e., C01, C02, and C03) were pooled and assembled independently in both MASH distance-based - and time-discrete assembly strategies.

Quality filtered forward and reverse FASTQ files of samples for the individual - and three co-assembly strategies were assembled using metaSPAdes v.3.13.1 and MEGAHIT v.1.2.9 with k-mere sizes 21, 33, 55, and 77. Following assembly and prior to binning, contigs of the MASH distance-based co-assemblies (*n* = 4), time-discrete co-assemblies (*n* = 12), and individual assemblies (*n* = 15) were pooled within each strategy, resulting in 6 pooled assemblies and 8 assemblies in total (6 pooled assemblies and two co-assemblies) that were used in downstream processing (Table S2). Contigs < 1kbp were filtered from all assemblies using seqtk (https://github.com/lh3/seqtk). This was followed by the removal of redundant contigs, i.e. duplicate and contained contigs, using the dedupe function of BBTools v38.76 (https://github.com/BioInfoTools/BBMap/blob/master/sh/dedupe.sh) for the pooled assemblies. QUAST v.5.0.2(46) was used to assess the quality of the processed assemblies with default parameters. Mapping rates were determined by mapping the quality-trimmed paired end reads to each assembly using BWA-MEM v0.7.17 (37) and then filtering unmapped reads using the view function of samtools v1.9 (38) with parameters: -hbS -F4. BAM files were subsequently sorted using the sort function of samtools v1.9 and then coverM v.0.4.0 (https://github.com/wwood/CoverM) was used to calculate contig-wise coverage with the method flag set to count. Prokka v1.14.6 (47) was used to identify coding DNA sequences (CDSs) in the contigs and to translate these CDSs to protein-coding amino acid sequences. Coding density was calculated by dividing the total CDS length (in Mbp) by the total assembly length (in Mbp). The blastp workflow of DIAMOND v.0.9.36 (48) was used to align the protein-coding amino acid sequences against the UniPort Knowledgebase (UniProtKB)/TrEMBL non-redundant (nr) protein database (https://www.uniprot.org/downloads) at an expected value (e-value) cutoff of 1 × 10^−3^ to identify high-scoring segment pairs (HSPs). Predicted protein-coding amino acid sequences that aligned with reference protein-coding amino acid sequences in the UniPort Knowledgebase (UniProtKB)/TrEMBL nr protein database were used to compute query/subject length ratios and query/subject length alignment ratios. These query/subject length and query/subject length alignment ratios were used as a measure to asses the extent of assembly fragmentation and misassembly.

Binning of contigs greater than 1.0kbp and 2.5kbp were performed using the analysis and visualization platform for ‘omics data (anvi’o) v6.1 (49). In this workflow, bins were generated using binning algorithms that combine tetranucleotide frequencies and coverage information across samples, including CONCOCT v.1.1.1 (13), MetaBAT v.2.12.1 (12), MaxBin v.2.2.4 (44). Since different binning tools reconstruct genomes at varying levels of completeness, a bin aggregation software, i.e., DAS Tool v.1.1.0 (45), was used to integrate the results of bin predictions made by CONCOCT, MetaBAT2 and MaxBin2 to optimize the selection on non-redundant, high-quality bin sets using default parameters. Bin statistics, including total size, number contigs, N50, GC content, etc., were obtained using the anvi-summarize function of anvi’o; while estimates of quality (completeness, redundancy, strain heterogeneity, etc.) were retrieved using the lineage-specific workflow of CheckM v.1.0.18 (50). Mapping rates were determined by mapping the quality-trimmed paired end reads to each bin using BWA-MEM v0.7.17 and then filtering unmapped reads using the view function of samtools v1.9 with parameters: -hbS -F4. BAM files were subsequently sorted using the sort function of samtools v1.9 and then coverM v.0.4.0 was used to calculate contig-wise coverage with the method flag set to count.

To further improve bin quality, individual bins with ≥ 50% completeness of a selected combination of assembler, assembly strategy, and binning approach were identified for reassembly (see results section). For this, properly paired quality-trimmed reads associated with individual bins of the selected assembly/binning approaches were extracted and stored into their FASTQ files using samtools v1.9 functions view and fastq, followed by assembly using metaSPAdes v.3.13.1 with k-mer sizes 21, 33, 55, and 77. The reassembled contigs were re-binned a second time using the appropriate original binning approach and bin statistics and mapping rates were determined as described above.

To obtain MAGs, bins were manually curated using the interactive interface of anvi’o v6.1. MAG characteristics and mapping rates were determined as described above. To assist in the identification of high- and medium-quality draft MAGs as define under the Minimum Information about a Metagenome-Assembled Genome (MIMAG) standards (51), ribosomal RNAs (rRNAs) and transfer RNAs (tRNAs) were detected with Prokka v. 1.14.6 (47). Pooled MAGs were dereplicated with dRep v2.6.2 (52) and clustered into species-level representative genomes (SRGs) at 95% average nucleotide identity (ANI). SRGs were classified using the classify workflow of the Genome Taxonomy Dataset Toolkit (GTDTk) v0.3.2 (53), which provides automated classification of bacterial genomes by placing them into domain-specific, concatenated protein reference trees. The phylogenomic workflow of anvi’o v6.1 was reproduced to construct a phylogenomic tree using a concatenated alignment of 37 single-copy ribosomal bacterial core genes.

### Statistical analysis

Statistical analysis was performed in R (40). Descriptive statistics and statistics on central tendency were performed using one-way analysis of variance (ANOVA) provided in the stats package. Significant ANOVA findings were further investigated by performing a post-hoc Tukey-Kramer test using the function Tukey.HSD with Bonferroni correction. Non-multidimensional scaling (NMDS) using Bray-Curtis and Jaccard dissimilarity indices were performed using metaMDS provided in the vegan package and permutational analysis of variance (PERMANOVA) were conducted using the function adonis of the vegan package. All plots were generated in R using ggplot2 (54).

## Results and discussion

### Summary of metagenomic sequencing of drinking water samples

On average 23.03 ± 9.57 ng DNA were extracted from the 1,500 ml filtered tap water samples harboring between 21.8 and 85.8 million cells (Table S1). A total of 1.05 billion (*M* ± *SD* = 87.67 ± 4.34 million reads) raw 150-nucleotide (nt) paired-end reads, ranging between 81.37 and 94.15 million reads per sample were generated from the DNA extracts of 12 samples, which had average 16S rRNA gene counts of 3.8×10^5^ ± 1.9×10^5^ copies/µl (Table S3). Control samples with average 16S rRNA gene counts of 4.1×10^1^ ± 8×10^0^ copies/µl had at least 3 × 10^2^-fold less raw paired-ended reads compared to samples (0.27 ± 0.12 million reads) (Table S3). Processing of the raw paired end reads following quality filtering and contaminant exclusion, removed on average 1.02 ± 0.23% of the reads per sample. The final sequence dataset consisted of 1.04 billion (86.75 ± 0.42 million) high-quality, processed reads with a lower and upper range of 80.51and 93.08 million reads per sample, respectively (Table S3). Nonpareil (39), was used to assess the coverage of sequencing effort (Table S4). The average coverage estimates across samples were 89.00 ± 3.00%, with a lower and upper range of 84.00 and 94.00%, respectively. This suggests that a sequencing depth of ∼81-94 million reads was sufficient to capture most of the microbial diversity in each sample.

### Evaluation of metagenome assembly quality for variable assembly strategy and metagenome assembler combinations

The performance of two de Bruijn graph-based assemblers, metaSPAdes and MEGAHIT that utilize iterative multiple k-mer approaches to improve assembly quality (43, 55) were assessed for three co-assembly strategies (i.e., co-assembly of all samples, MASH distance-based assembly and time-discrete assembly) and assembly of individual samples (Table S2). Inclusion of various assembly strategies allow for the assessment of assembly performance in terms of computational requirements (i.e., RAM usage, assembly runtime per processing core, etc.) and assembly quality at varying levels of diversity as well as sequence depth and coverage. The metaSPAdes assemblies required more computing recourses, i.e., demanded higher memory limits and threads, and had runtimes that were up to 6-fold longer when compared to the MEGAHIT assemblies (Table S5), which confirms previous findings (18, 19, 56). As expected, co-assembly strategies (i.e., co-assembly of all samples, MASH distance-based assembly and time-discrete assembly) for both metaSPAdes and MEGAHIT were associated with longer runtimes (Table S5). Amongst the metaSPAdes assemblies, time-discrete assembly had the longest runtime (195 hours summed across all assemblies), followed by co-assembly (100 hours) and MASH-distance based assembly (28 hours summed across all assemblies). Similar observations were made for the MEGAHIT assemblies (Table S5).

Evaluating *de novo* assembly quality for environmental samples is challenging due the lack of a ground truth reference assembly for comparison (57). As a result, we used measures of contiguity (i.e., total assembly size, maximum contig length, N50, L50, etc.), gene calling and quality (i.e., coding DNA sequence (CDS), coding density), mapping rate, and rate of gene fragmentation and misassembly to assess the quality of the assemblies (Table S5). In total, between 9,959,586 and 114,386,414 contigs were generated across the metaSPAdes and MEGAHIT assemblies. Most of the assembled contigs ∼98.39% had lengths below 1kbp and were not used in downstream analysis. Duplicate and contained contigs that accounted for between 10 and 42% of the filtered contigs (>1kbp) were found amongst the MASH distance-based - and time-discrete co-assemblies, and individual assemblies of metaSPAdes and MEGAHIT and removed (Table S5). Duplicate contigs were define as contigs sharing 100% sequence similarity over the entire length, while contained contigs included shorter contigs that were 100% similar to a longer contig over their length. The average total number contigs per assembly strategy kept after removing contigs shorter than 1kbp and redundant contigs were 506,898 (*SD* = 164,157). For each assembly strategy, the metaSPAdes assemblies produced between 10 and 20% more contigs greater than 1kbp compared to the number of contigs that were generated from the MEGAHIT assemblies (Table S5). Differences between assemblies were more apparent when metrics related to assembly contiguity were compared. Irrespective of the assembly strategy, the total assembly length of the metaSPAdes assemblies were greater when compared to the assemblies of MEGAHIT (Figure 1A). MetaSPAdes time-discrete assembly had the greatest assembly length (2,940.15 Mbp), followed by individual assembly (2,037.97 Mbp), co-assembly (1,488.27 Mbp), and MASH distance-based assembly (1,147.06 Mbp). Since larger assembly lengths are not always indicative of better assembly quality (56), N50 estimates representing a weighted medium contig size were considered. The metaSPAdes assemblies generated contigs with higher N50 estimates when compared to the MEGAHIT assemblies (Figure 1B). Time-discrete assembly of metaSPAdes had the highest N50 estimates (6.77 kbp), followed by individual assembly (5.67 kbp), MASH distance-based assembly (5.65 kbp), and co-assembly (5.32 kbp). The higher N50 estimates of the metaSPAdes assemblies indicate that these assemblies contain a lower proportion of small contigs and therefore are less fragmented assemblies (1, 56, 58, 59).

**Figure 1.**
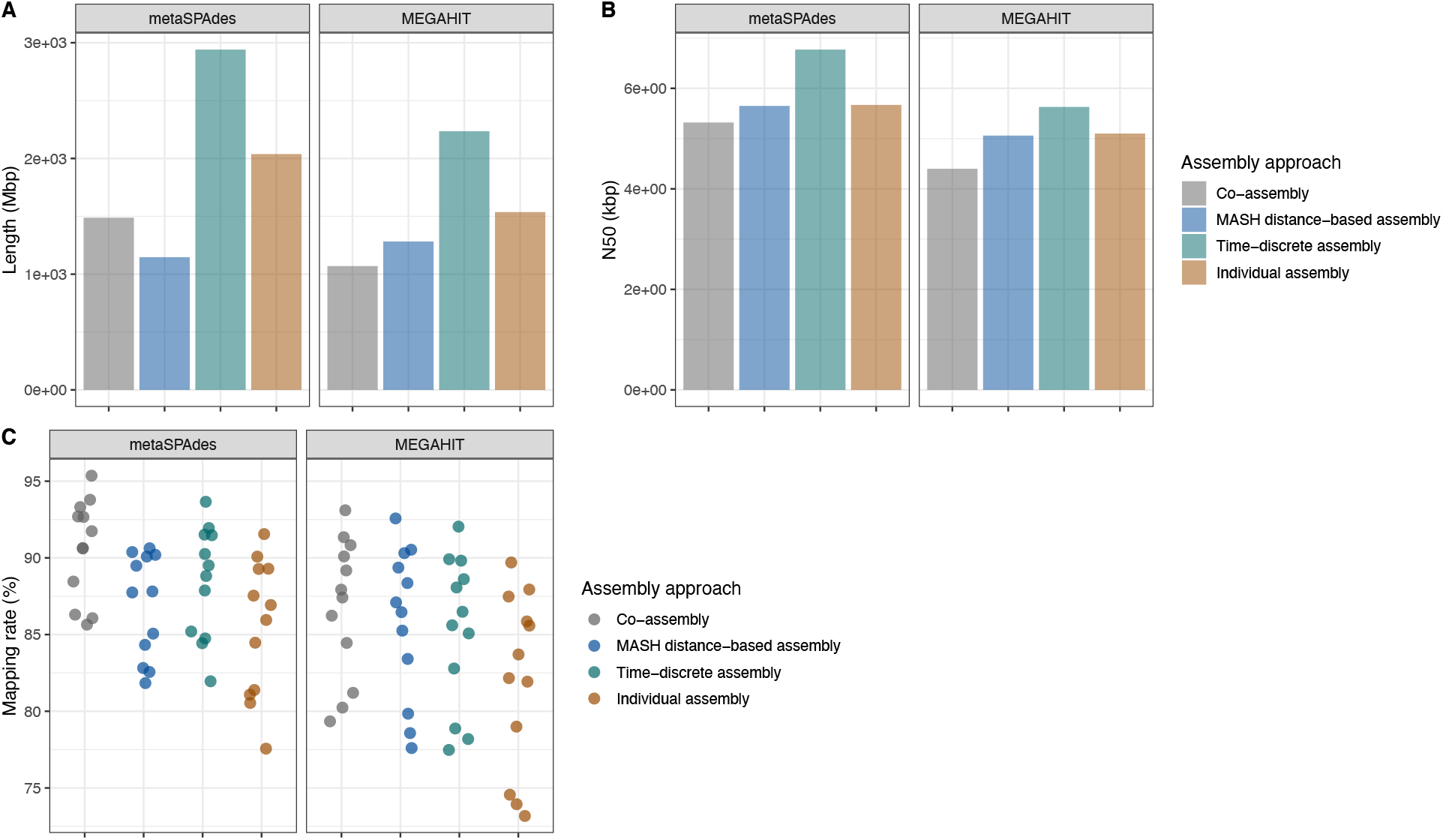
Comparison of assembly characteristics associated with 4 different assembly strategies (co-assembly (grey), MASH distance-based assembly (blue), time-discrete assembly (green) and individual assembly (orange)) that were assembled with metaSPAdes and MEGAHIT. Assembly characteristics were determined using non-redundant contigs larger than 1kbp of co-assemblies and pooled MASH distance-based, time-discrete, and individual assemblies. Assembly characteristics included: **A)** Total assembly length; **B)** N50 estimates; and **C)** Proportion reads of 12 drinking water samples (•) that were mapped against the non-redundant, filtered assemblies. For a complete list of estimates, please refer to Tables S5.

Although the metaSPAdes assemblies were associated with 10-30% more CDSs compared to the MEGAHIT assemblies, the coding densities were similar across the assemblies of metaSPAdes and MEGAHIT, and range between 0.77 and 0.80 (Table S5). CDSs were blasted against the UniProtKB/TrEMBL nr protein database to identify HSPs, i.e., sequence pairs sharing high alignment scores, at an expected value (e-value) cutoff of 1 × 10^−3^. Across the assembly strategies, between 71 and 76% of the CDSs shared a high degree of similarity against the reference amino acid sequences in the UniProtKB/TrEMBL nr protein database and had average e-values and bit scores of 3.09 × 10^−6^ ± 3.91 × 10^−5^ and 387.00 ± 305.65, respectively (Table S5). Assembly fragmentation was assessed by analyzing the ratio between the lengths of CDSs (query length (qlen)) and their top hits in the UniProtKB/TrEMBL nr protein database (subject length (slen)), with lower qlen:slen ratios indicating less gene fragmentation and thus lower assembly fragmentation. Similar distributions in qlen:slen ratios were observed across the assemblies of metaSPAdes and MEGAHIT, with only between 30 and 36% of the CDSs having qlen:slen ratios ranging between 0.95 and 1 (Figure S3A, Table S6.1). This suggests that the vast majority of CDSs across both assemblers and all assembly strategies were likely fragmented.

The CDSs of the metaSPAdes assemblies were less fragmented than those from the MEGAHIT assemblies; hence had higher mean qlen:slen ratios (Table S6.1, Figure S3B). Post-hoc comparisons using the Tukey HSD test indicated statistically significant differences between all metaSPAdes and MEGAHIT assemblies (Tukey HSD test, all *p* < 0.05) (Tables S6.2, S6.3). metaSPAdes time-discrete assembly had the greatest mean qlen:slen ratio (0.901 ± 0.307), followed by MASH distance-based assembly (0.895 ± 0.315), individual assembly (0.895 ± 0.306), and co-assembly (0.892 ± 0.320). Similar observations were made for the MEGAHIT assembly strategies (Table S6.1). Though significant differences were found, the effect sizes of these differences was small (effect size (η^2^) = 1.15E-04 and 2.39E-04 for metaSPAdes and MEGAHIT, respectively), suggesting that only 0.01 and 0.02% of the change in qlen:slen ratios can be accounted for by the assembly strategy for metaSPAdes and MEGAHIT, respectivly. Statistically significant differences in the variation around the mean qlen:slen ratios were observed for the assemblies of metaSPAdes (coefficient of variation (C_v_) = 0.35 ± 0.01) and MEGAHIT (C_v_ = 0.36 ± 0.01) (signed-likelihood ratio test (SLRT) and asymptomatic test, all *p* < 0.05) (Table S6.4). Associations between the mean qlen:slen ratios and C_v_ estimates indicated that the metaSPAdes assembly strategies were associated with higher qlen:slen ratios and lower C_v_ estimates compared to the assemblies of MEGAHIT (Figure S3B). The ratio between the alignment lengths of CDSs (query alignment length (qalignlen)) and their top hits in the UniProtKB/TrEMBL nr protein database (subject alignment length (salignlen)) were used to evaluate potential misassembly due to the presence of insertion-deletion (indels) in genes (Table S7.1). Similar distributions in qalignlen:salignlen ratios were observed across the assemblies of metaSPAdes and MEGAHIT, with between 59 and 66% of the CDSs having qalignlen:salignlen ratios that ranged between 0.95 and 1 (Figure S3C, Table S7.1). Though statistically significant differences in the mean qalignlen:salignlen ratios were observed for the assemblies of metaSPAdes and MEGAHIT (ANOVA, all *p* < 0.05) (Table S7.2, S7.3), the effect size of the differences were small (η^2^ = 6.22 × 10^−5^ and 3.74 × 10^−5^ for metaSPAdes and MEGAHIT, respectively). Similarly, small but statistically significant differences in the variation around the mean qalignlen:salignlen ratios of the metaSPAdes and MEGAHIT assembly strategies were observed (C_v_ range = 0.02 – 0.03 for metaSPAdes and MEGAHIT assemblies) (SLRT and asymptomatic test, all *p* < 0.05) (Table S7.4). Overall, these results suggests that while metaSPAdes results in significantly less fragmented assemblies with lower rates of genes fragmentation, the effect size of this difference on CDS quality is small.

The proportion of sequencing information retained following assembly was determined by mapping the quality-trimmed paired end reads of each sample to the non-redundant, filtered metaSPAdes and MEGAHIT assemblies. Although no statistically significant differences between the mean read mapping rates of corresponding metaSPAdes and MEGAHIT assembly strategies were observed (Tukey HSD, all *p* < 0.05) (Table S8), the metaSPAdes assemblies had mean mapping rates that were higher when compared to the MEGAHIT assemblies (Figure 1C, Table S5). Amongst the metaSPAdes assemblies, co-assembly of all samples had the highest mapping rate (90.61 ± 3.28%), followed by time-discrete assembly (88.45 ± 3.64%), MASH distance-based assembly (86.91 ± 3.39%) and individual assembly (85.47 ± 4.45%). Similar observations were made for the MEGAHIT assemblies. Overall, the metaSPAdes assemblies had larger and more contiguous assemblies with read mapping rates > 85%. This confirms previous findings (1, 19, 59, 60).

### Evaluation of binning results for combination of assembly strategies, assemblers, and binning approaches

Unrefined bin sets were generated from the metaSPAdes and MEGAHIT assemblies of each assembly strategy using original binning algorithms that combine tetranucleotide frequencies and coverage information across samples (12, 13, 44), i.e., CONCOCT v.1.1.1, MetaBAT v.2.12.1 and MaxBin v.2.2.4 as well as DAS Tool v.1.1.0 that integrates results of bin predictions made by original binning algorithms to optimize the selection on non-redundant, high-quality bin sets (45). This resulted in 64 assembly/binning combinations (*n* = 32 assembly/binning combinations for bin sets that were constructed using larger than 1kbp and 2.5kbp contigs, respectively) (Table S9). MASH distance based NMDS clustering of the unrefined bin sets indicated that the bin sets clustered based on assembly/binning approach rather than contig size cutoff used for binning (Figure S4). The importance of assembly/binning approach as compared to contig size cutoff was further confirmed by PERMANOVA analyses (Table S10). The minimum contig threshold (1kbp or 2.5kbp) that were selected for binning, explained a smaller proportion of the variation ∼2% (PERMANOVA, *F*(1) = 36.16, *R*^*2*^ = 0.02, *p* < 0.05) as compared to 96% of the variation that was explained by the assembly/binning approach choice (PERMANOVA, *F*(31) = 46.24, *R*^*2*^ = 0.96, *p* < 0.05). However, the unrefined bin sets that were generated using contigs > 1kbp produced about 20% more unrefined bins with ≥ 50% completeness, when compared to the unrefined bin sets that were generated using contigs > 2.5kbp (Figure 2, Table S9). These unrefined bins were furthermore associated with mapping rates that were between 5 and 20% higher when compared to the unrefined bins that were generated using contigs > 2.5kbp (Figure 2, Table S9). These finding suggest that binning with contigs > 1kbp allows for a more accurate representation of the microbial diversity.

**Figure 2.**
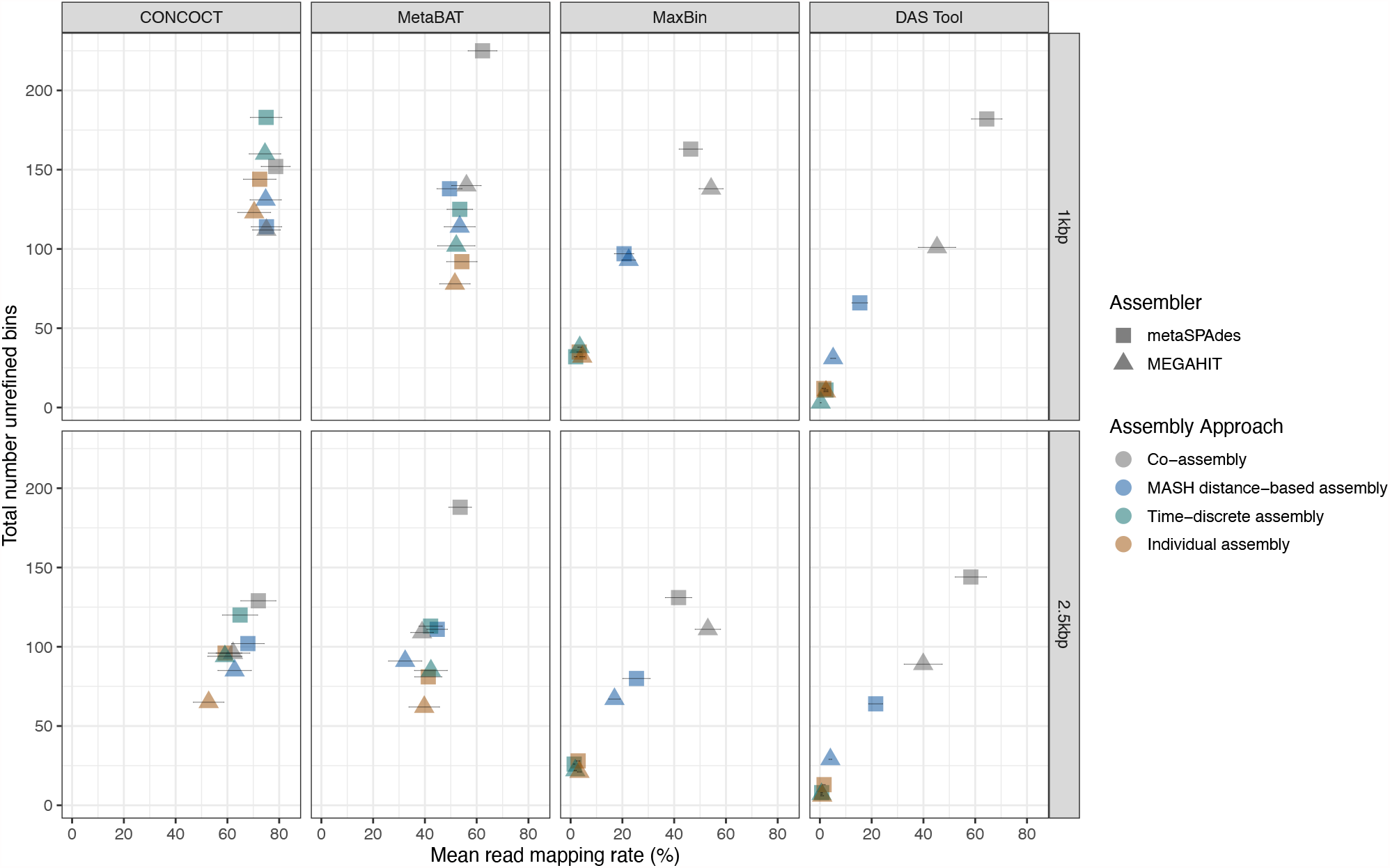
Association between total number bins and mean read mapping rates of sample reads mapped against the unrefined bins with completeness ≥ 50% that were assembled with different assembly approaches (co-assembly (grey), MASH distance-based assembly (blue), time-discrete assembly (green) and individual assembly (orange)) using metaSPAdes (▲) and MEGAHIT (▪) and binned with CONCOCT, MetaBAT2, MaxBin2 and DAS Tool. Error bars indicate standard errors of read mapping rates.

### Marked improvements in bin qualities following re-assembly and curation

Across the 64 assembly/binning combinations, greater than 1kbp and 2.5kbp contig size bin sets of 4 assembly/binning approaches; hence, 8 assembly/binning combinations in total, consistently produced the highest number bins (completeness ≥ 50%) and mapping rates greater than 50%. These assembly/binning approaches included co-assembly strategies of metaSPAdes: (i) metaSPAdes co-assembly + CONCOCT, (ii) metaSPAdes time-discrete assembly + CONCOCT, (iii) metaSPAdes co-assembly + MetaBAT2, and (iv) metaSPAdes co-assembly + DAS Tool (Figure 2). The unrefined CONCOCT bin sets of the co-assembled metaSPAdes assemblies (i.e., metaSPAdes co-assembly + CONCOCT and metaSPAdes time-discrete-assembly + CONCOCT), consisted of fewer bins and bins that were significantly greater in size (Table S11). In particular, for metaSPAdes co-assembly + CONCOCT the average bin size were 8.15 ± 8.12 Mbp and 6.41 ± 5.26 Mbp for 1kbp and 2.5kbp constructed bins, while metaSPAdes time-discrete-assembly + CONCOCT had average bin sizes of 14.05 ± 15.06 Mbp (contigs > 1kbp) and 15.01 ± 12.59 Mbp (contigs > 2.5kbp). These bins were also associated with large redundancy estimates that averaged above 60% and average strain heterogeneity estimates greater than 20% (Figure 3). These findings suggest that the unrefined CONCOCT bin sets likely consist of multi-genome or chimeric bins and highlighted the need for reassembly of individual bins and/or bin curation (61). The remaining co-assembly strategies of metaSPAdes, i.e., metaSPAdes co-assembly + MetaBAT2 and metaSPAdes co-assembly + DAS Tool, generated more bins and bins with lower redundancy estimates that average below 15% (Figure 3). To improve the quality of the unrefined bin sets, bins with greater than 50% completeness of the 8 assembly/binning combinations were independently subjected to reassembly. Proper paired quality-trimmed reads associated with the bins were extracted, converted to FASTQ format, and then reassembled using metaSPAdes and re-binned using the appropriate original binning approach. Following reassembly, the reassembled unrefined bin sets of metaSPAdes co-assembly + CONCOCT and metaSPAdes time-discrete assembly + CONCOCT consisted of bins that were notably smaller in size (Table 11). Specifically, the metaSPAdes time-discrete + CONCOCT reassembled average unrefined bin sizes of the 1kbp and 2.5kbp bins (4.21 ± 2.75 Mbp and 3.97 ± 1.66 Mbp, respectivly) where at least 4-fold lower when compared to the original unrefined bin sizes. Similar observations were made for metaSPAdes co-assembly + CONCOCT that had an average 2-fold reduction in bin size (Table 11). Furthermore, reductions in bin size across the co-assembled CONCOCT bins were accompanied by improvements in bin quality (Figure 3). These improvements were associated with reduced redundancy estimates across the 1kbp and 2.5kbp reassembled unrefined bin sets of metaSPAdes co-assembly + CONCOCT (26.91 ± 47.05% and 23.86 ± 42.87%) and metaSPAdes time-discrete assembly + CONCOCT (21.52 ± 41.98% and 11.08 ± 27.92%). These findings suggest that the unrefined CONCOCT bins set consisted of chimeric bins that were resolved with reassembly. This improvement in bin quality after reassembly is consistent with previous findings (61). In contrast, no improvements in bin quality were observed in the reassembled bin sets of MetaBAT2 and DAS Tool (Figure 3). Specifically, the reassembled unrefined 1kbp and 2.5kbp bin sets of metaSPAdes co-assembly + MetaBAT2 maintained smaller bin sizes (4.16 ± 3.04% and 3.42 ± 2.42%) as well as average redundancy estimates below approximately 10% (11.05 ± 29.38% and 4.41 ± 11.71%) follow reassembly. Similar observations were made for metaSPAdes co-assembly + DAS Tool. This was expected as higher quality bins were associated with both MetaBAT and DAS Tool bin sets prior to reassembly.

**Figure 3.**
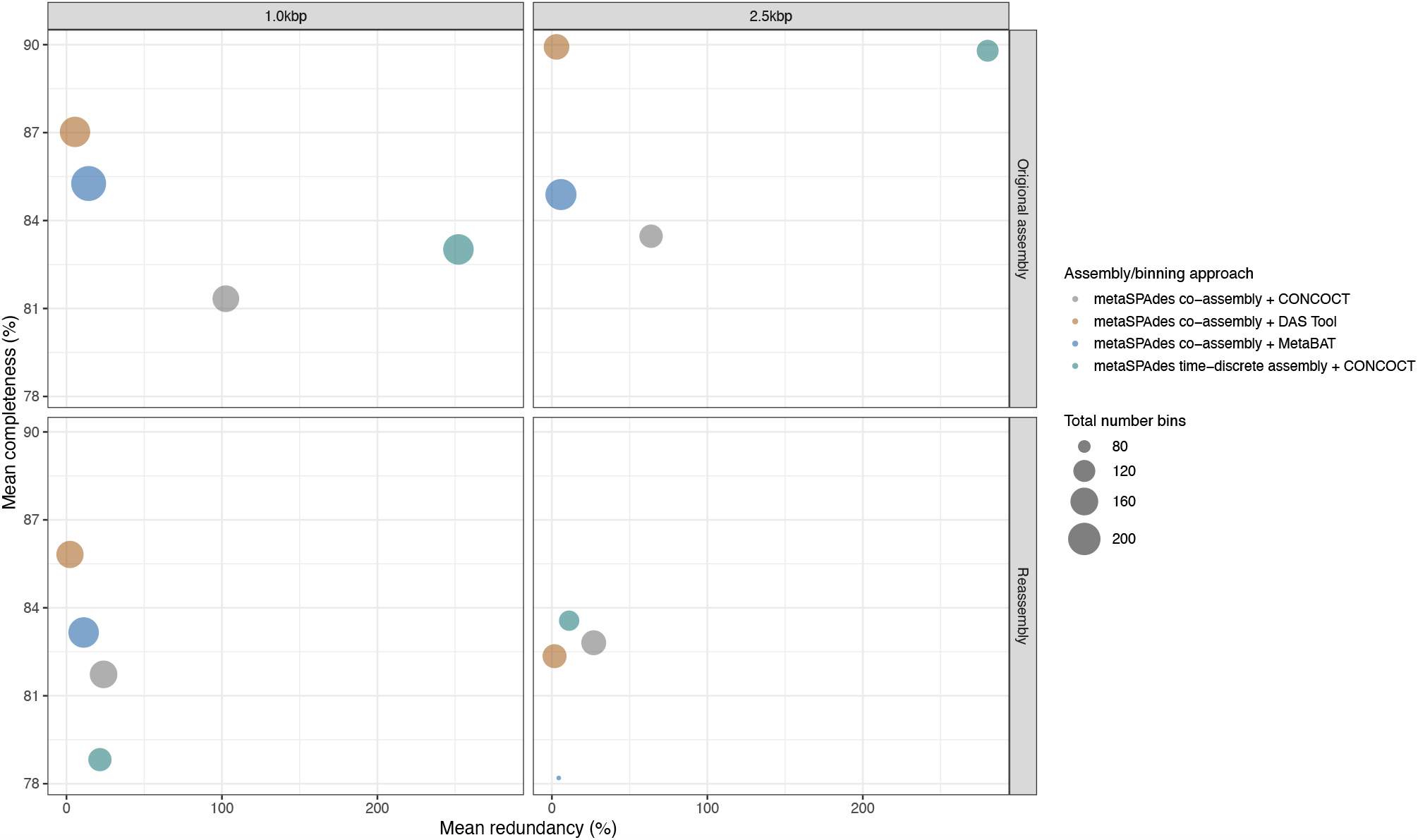
Bubble plot showing the total number of bins (depicted by size) with mean completeness and redundancy estimates of unrefined bin sets (completeness estimates > 50%) that were generated using 4 assembly/binning approaches (metaSPAdes co-assembly + CONCOCT (orange), metaSPAdes co-assembly + DAS Tool (grey), metaSPAdes co-assembly + MetaBAT2 (blue) and metaSPAdes time-discrete assembly + CONCOCT (green)) and binning of contigs greater than 1kbp and 2.5kbp.

### Metagenome assembled genomes shared across assembly/binning approaches

The reassembled CONCOCT bin sets (i.e., metaSPAdes co-assembly + CONCOCT and metaSPAdes time-discrete assembly + CONCOCT) and original assembled MetaBAT and DAS Tool bin sets were manually curated using the interactive interface of anvi’o v6.1 (49) to obtain final MAGs (completeness ≥ 50% and redundancy < 10%). In total 1,279 MAGs were generated across the 4 assembly/binning approaches that were constructed using contigs > 1kbp (*n* = 673) and contigs > 2.5kbp (*n* = 606). Approximately 98% (*n* = 1,259) of the MAGs that were identified met the MIMAG strandard (51) for medium-quality draft genomes, while only 20 MAGs were classified as high-quality draft genomes (Table S12.1). The limited number high-quality MAGs were mainly due to the absence of full complement of rRNA genes. Depending on the assembly/binning approach, between 75 and 83% of the MAGs lacked 16S rRNA genes, while between 10 and 16% of the MAGs consisted of fragmented 16S rRNA gene(s) (Table S12.1). MAGs often lack 16S rRNA genes due to their conserved and repetitive nature, which results in fragmented assemblies (22, 62, 63). Overall, none of the assembly/binning strategies produced sufficient high-quality MAGs as define under MIMAG standards. Alternative sequencing technologies (e.g., long-read) may be able to successfully reconstruct full complement rRNA genes to increase the number high-quality MAGs (5).

The MAGs across the 4 assembly/binning approaches shared similar characteristics in terms of contiguity (i.e., total length and N50) and quality (i.e., completeness, redundancy, and strain heterogeneity) (Figure 4A and Table S12.1). Overall, the curated MAG sets of the assembly/binning approaches retained more than 66% of the sequencing information (Table S12.1). The curated MAG sets of DAS Tool and MetaBAT had higher mapping rates ∼70%, when compared to the curated MAG sets of CONCOCT. As shown in Figure 4B, MAGs that were reconstructed using a minimum contig length of 1kbp had slightly higher mapping rates, when compared to the mapping rates of the MAGs that were reconstructed using a minimum contig length of 2.5kbp. These differences in read mapping rates were not statistically significantly different between corresponding assembly/binning strategies that used minimum contig threshold of 1kbp and 2.5kbp for MAG reconstruction, respectively (Tukey HSD test, all *p* > 0.05) (Table S12.2).

**Figure 4.**
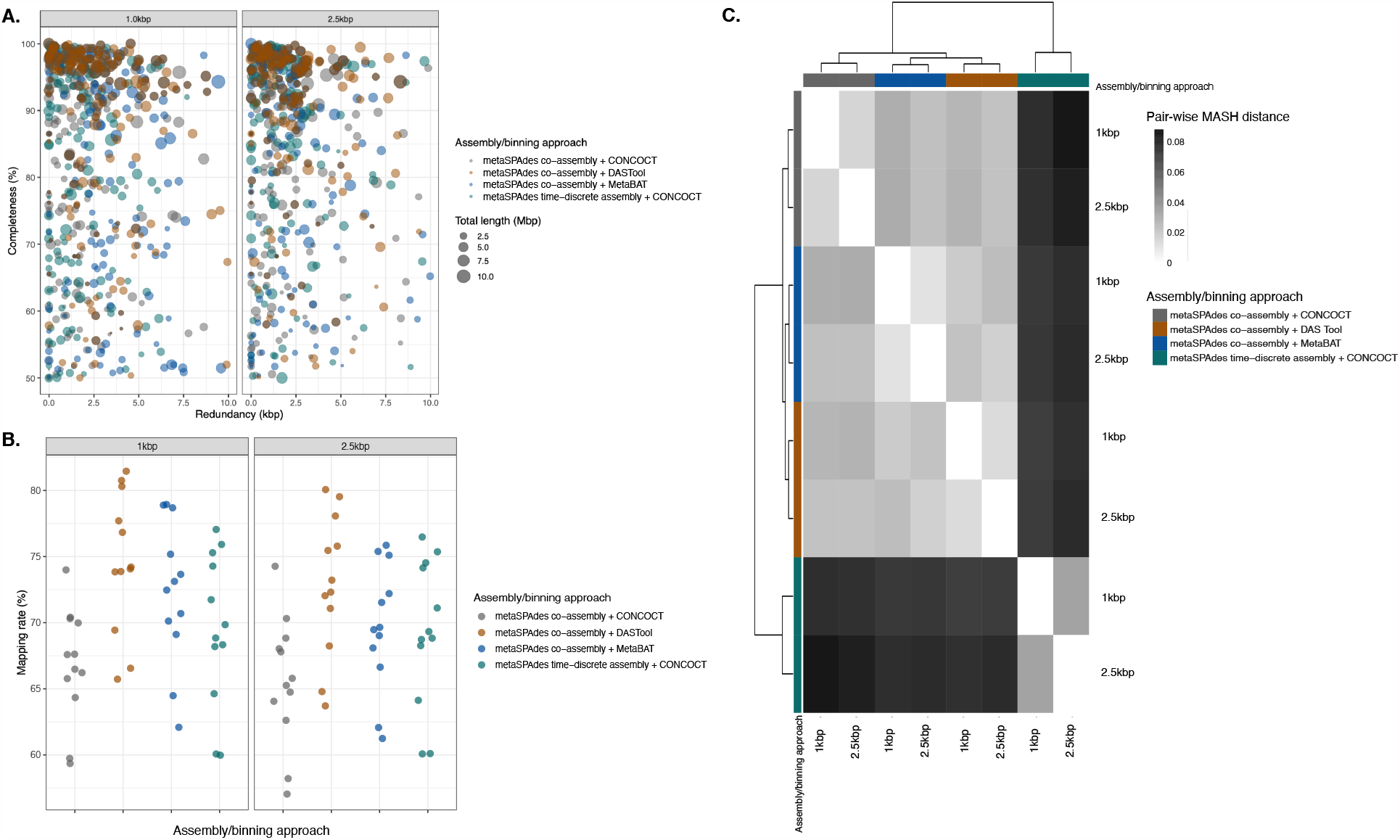
Summary statistics and characteristics of 1,279 curated MAGs that were generated across the 4 assembly/binning approaches (metaSPAdes co-assembly + CONCOCT (grey), metaSPAdes co-assembly + DAS Tool (orange), metaSPAdes co-assembly + MetaBAT2 (blue) and metaSPAdes time-discrete assembly + CONCOCT (green)) using contigs larger than 1kbp (*n* = 673) and 2.5kbp (*n* = 606), respectively. **A)** Bubble plot showing total MAG size (depicted by size) and completeness (x-axis) and redundancy (y-axis) estimates 1279 curated MAGs that were generated for each of the 4 assembly/binning approaches. **B)** Proportion reads of 12 drinking water samples (•) that were mapped against the curated MAGs of each assembly/binning approach. **C)** Comparison of the curated MAGs nucleotide composition across the different assembly/binning approaches according to MASH distance. The heatmap are colored according to MASH distance; white denotes a distance of 0. Labels on the x- and y-axis are colored according to assembly/binning approach and clustering is done using Euclidean distance. For a complete list of continuity and quality estimates, please refer to Table S12.

The curated MAG sets clustered by assembly/binning approach based on MASH distance estimates that explained approximately 91% of the variation in the nucleotide composition (PERMANOVA, *F*(3) = 16.80, *R*^*2*^ = 0.91, *p* < 0.05) (Figure 4C, Table S13). Though the minimum contig threshold (1kbp or 2.5kbp) that were selected for binning explained a smaller proportion of the variation ∼3% this was not significant (PERMANOVA, *F*(1) = 1.73, *R*^*2*^ = 0.03, *p* > 0.05) (Table S13). Based on MASH distance estimates, the differences in nucleotide composition of the curated MAGs between assembly/binning approaches were small, ranging between 0.005 and 0.08. metaSPAdes time-discrete assembly clustered separately from the other assembly/binning strategies, suggesting differences in the nucleotide composition of these curated MAGs.

Similarities in the nucleotide composition of the curated MAG sets and comparable MAG characteristics (i.e., continuity and quality) suggests the presence of overlapping MAGs across the assembly/binning approaches that likely represents the same species. The presence of overlapping MAGs across the assembly/binning approaches were investigated by aggregating all the MAGs and then clustering them using a 95% ANI threshold to identify species-level representative genomes (SRGs). Although the species concept for prokaryotes is controversial, this operational definition is commonly used and considered a golden standard (64, 65). A total of 233 SRGs with average dRep quality scores (calculated as: A*Completeness - B*Contamination + C*(Contamination * (strain_heterogeneity/100)) + D*log(N50) + E*log(size) + F*(centrality - S_ani)) (52) of 74.40 ± 21.80% were identified across the assembly/binning approaches (Table S14). These SRGs had average sizes of 3.46 ± 1.72% and were near complete (81.98 ± 16.39%) with redundancy estimates less than 10%. Taxonomic classification of the SRGs using GTDB-Tk classified 33 SRGs to species level, 178 to genus level, 217 to family level, and 233 to order, class, and phylum level (Figure 5A and Table S14).

**Figure 5.**
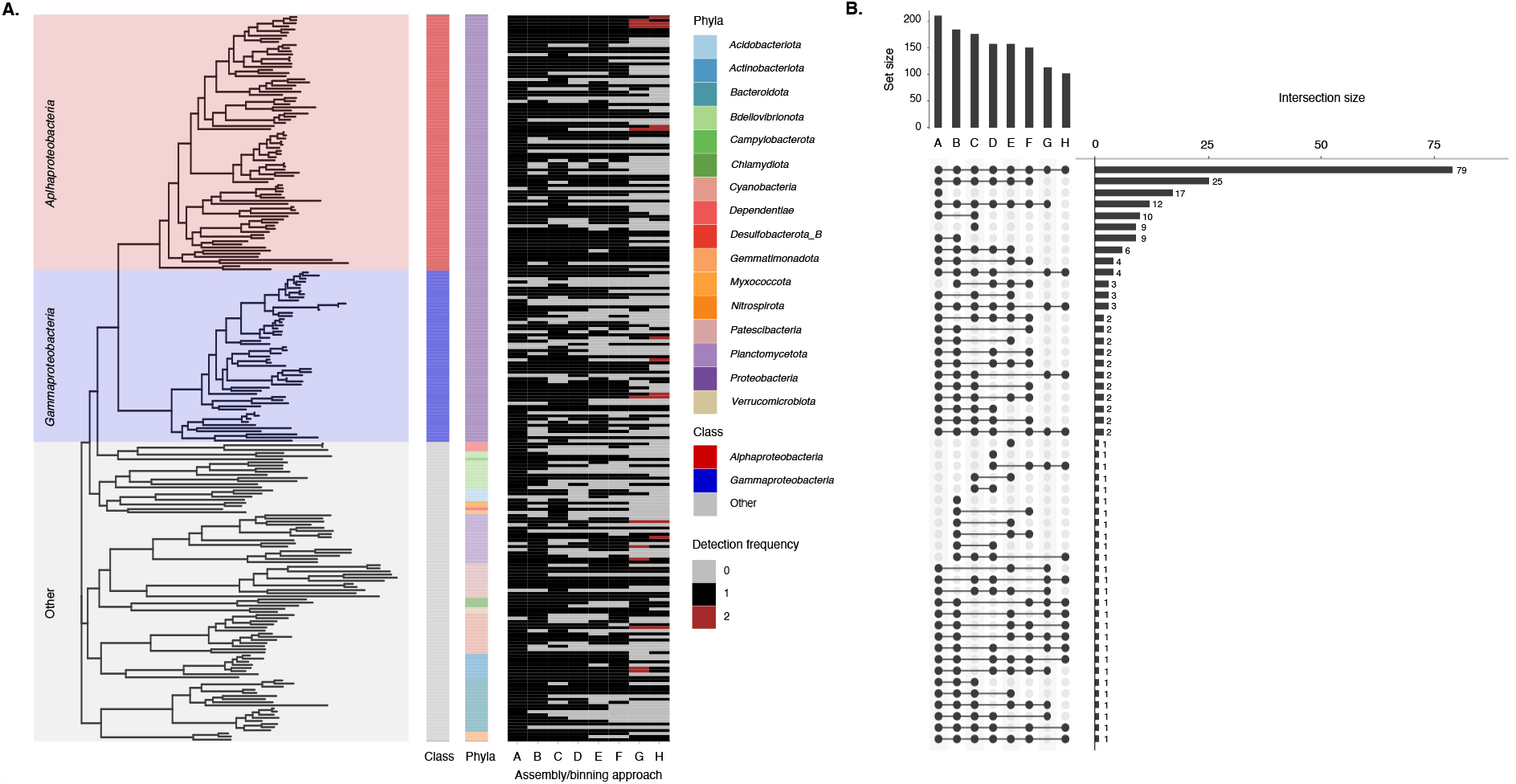
**A)** Phylogenomic analysis of 233 near-complete SRGs inferred from 37 single-copy ribosomal bacterial core genes. The two inner panels represent taxonomic classification of 16 bacterial phyla and class-level classification of dominating bacterial phyla, *Proteobacteria* representing *Alphaproteobacteria* (red) and *Gammaproteobacteria* (blue). The outer panel represents a presence/absence summary plot showing the frequency distribution of MAGs that were constructed using contigs greater than 1kbp and 2.5kbp across 4 assembly/binning approaches. Grey denotes absence, while black and red denotes the presence of a singular MAG or duplicate MAGs that demonstrated ≥ 95% ANI, respectively. **B)** UpSetR plot showing the distribution of species-level representative genomes (SRGs) that demonstrated ≥ 95% ANI between the 4 assembly/binning approaches in which MAGs were constructed using contigs greater than 1kbp and 2.5kbp. For the assembly/binning approaches: A = metaSPAdes co-assembly + MetaBAT2 (1kbp), B = metaSPAdes co-assembly + MetaBAT2 (2.5kbp), C = metaSPAdes co-assembly + DAS Tool (1kbp), D = metaSPAdes co-assembly + DAS Tool (2.5kbp), E = metaSPAdes co-assembly + CONCOCT (1kbp), F = metaSPAdes co-assembly + CONCOCT (2.5kbp), G = metaSPAdes time-discrete assembly + CONCOCT (1kbp), and H = metaSPAdes time-discrete assembly + CONCOCT (2.5kbp)).

Approximately 34% (*n* = 79) of the SRG were shared across the assembly/binning approaches where they accounted for between 39 and 48% of the sequencing data (Figure 5B and Table S15). These SRG had better quality with average completeness and redundancy estimates of 94.03 ± 8.3% and 1.3 ± 1.17%, respectively. Unique SRGs represented 10% (n = 29) of the total SRGs, while the largest proportion of the SRGs ∼ 43% (*n* = 125) where shared between two or more assembly/binning approaches (but not all) (Figure 5B). The latter shared similar quality characteristics, compared to the SRG that were shared across all the assembly/binning approaches and accounted for between 18 and 32% of the sequencing data. Overall, metaSPAdes co-assembly + MetaBAT2 (with contigs > 1kbp) retained more SRGs (*n* > 200) and where able to reconstruct MAGs that where not detected in the other assembly/binning approaches (Figure 5A). Though metaSPAdes time-discrete assembly + CONCOCT where associated low number SRGs (*n* < 120), 12 duplicate SRGs sharing > 95% ANI were identified within the 1kbp and 2.5kbp approaches, respectively. These SRGs were likely sub-species, suggesting that the metaSPAdes time-discrete assembly + CONCOCT assembly/binning approach can differentiate between closely related species that were, otherwise collapsed or considered a singular strain using the other assembly/binning approaches. This highlights the potential for utilizing multiple approaches for not just binning, but also assembly strategies as this can assist in the recovery of a greater proportion of populations in metagenomes.

Reads from all samples were mapped to the 233 SRGs and their relative abundance in each sample for all assembly/binning and contig size strategy was estimated based on the presence/absence of the bin in the respective strategy. Variability in the microbial community structure and membership between the assembly/binning approaches was visualized by ordinating the samples in multidimensional space. As shown in Figure 6, the samples cluster by time point which explained approximately 64% of the variation in the community membership (PERMANOVA, *F*(11) = 108.88, *R*^*2*^ = 0.637, *p* < .05) and 70% of the variation in community structure (PERMANOVA, *F*(11) = 50.92, *R*^*2*^ = 0.703, *p* < .05) (Table S16.1, S16.2). The remaining variables, i.e., assembly/binning approach and contig size, accounted for smaller but significant proportion of the variation. In particular, assembly/binning approach explained about 29% of the variation in community membership (PERMANOVA, *F*(3) = 182.78, *R*^*2*^ = 0.291, *p* < .05), while contig size explained 3% (PERMANOVA, *F*(3) = 55.50, *R*^*2*^ = 0.03, *p* < .05). Similar observations were made for structure-based analysis (Table S16.1). Although no clustering by assembly/binning approach and contig size were observed, the average dissimilarity in community structure and membership between time points were about 37 and 53%, respectively (d_BC_ and d_J_ = 0.369 ± 0.11 and 0.528 ± 0.13). These findings suggest that while temporal dynamics of the drinking water microbiome are largely retained despite variation in genome-centric metagenomic workflows, the choice of assembly/binning strategy, in particular, can have a significant impact on the structure and membership of the drinking water microbiome and should not be overlooked.

**Figure 6.**
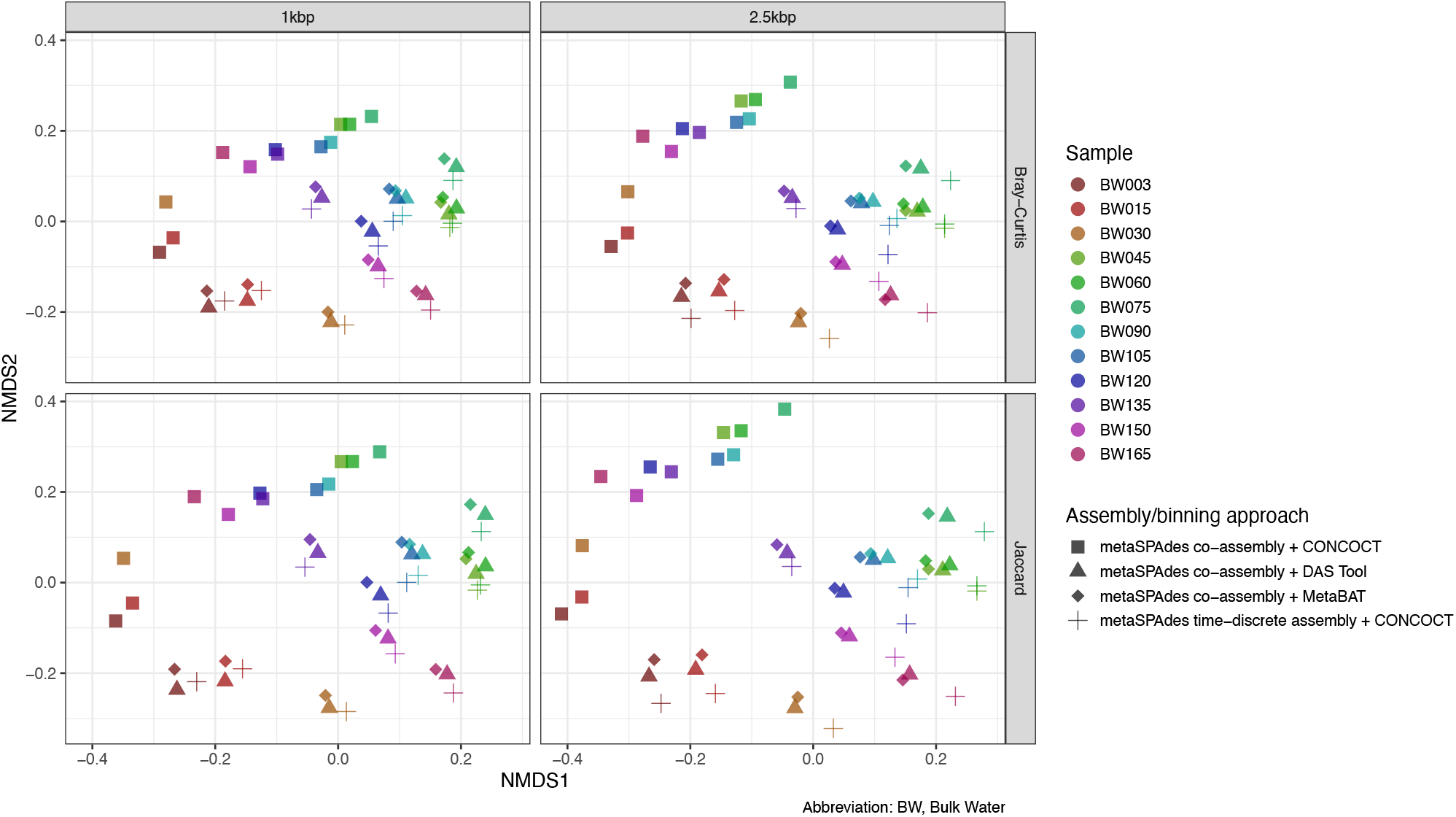
Structure-based non-metric multidimensional scaling (NMDS) of Bray-Curtis dissimilarity matrices and membership-based Jaccard distances as inferred using the abundance information (in RPKM) of all MAGs identified across the assembly/binning approaches. Points represents samples (or time points) (see supplementary Table S1), and shapes represents the assembly/binning approaches, i.e., metaSPAdes co-assembly + CONCOCT (▪), metaSPAdes co-assembly + DAS Tool (▲), metaSPAdes co-assembly + and MetaBAT2 (♦) and metaSPAdes time-discrete assembly + CONCOCT (+)).

## Conclusion

This study evaluated the performance of a combination of *de novo* assembly strategies and binning algorithms for time-series metagenomic data for drinking water microbial communities, to identify an ideal combination of assembly and binning approaches that allows for the generation of high quality metagenomic assemblies and MAGs. Overall, metaSPAdes co-assembly strategies, i.e., co-assembly of all samples and time discrete assembly, produced less fragmented and larger assemblies that retained the maximum amount of metagenomic information. Re-assembly and binning followed by manual curation significantly improved MAG qualities in situation with unresolved multi-genome or chimeric bins. Though none of the assembly/binning strategies were able to reconstruct high-quality MAGs due the absence of a full complement of rRNA genes, metaSPAdes co-assembly + MetaBAT2 retained the highest number medium-quality MAGs and where able to reconstruct MAGs that where not detected with the other assembly/binning approaches. Moreover, metaSPAdes time-discrete assembly + CONCOCT where able to differentiate between closely related species that were, otherwise collapsed or considered a singular strain using the other two assembly/binning approaches. Our study also finds that the choice of assembly/binning strategy can have a significant impact on the membership and structure of the microbial community as inferred from presence/absence and relative abundance of MAGs. This combined with the fact that a significant proportion of SRG’s were not reconstructed using any single approach, highlights the need to utilize multiple assembly/binning approaches for MAG recovery. We therefore recommend utilizing multiple assembly, binning and binning aggregating strategies followed by dereplication to maximize the recovery of non-redundant MAGs that may more fully represent the microbial populations in drinking water samples.

## Data availability

Raw sequence reads and 233 SRGs are available on NCBI at Bioproject number PRJNA745168 and PRJNA745370, respectivly. The 8 assemblies and 1,279 curated MAGs are available on datadryad at https://doi.org/10.5061/dryad.ksn02v74q.

## Funding sources

This research was supported by NSF Award number: 1749530.

## Notes

### Competing Interest Statement

The authors have declared no competing interest.

