## Supplementary material for "Evaluating *de novo* assembly and binning strategies for time-series drinking water metagenomes": Figure S4

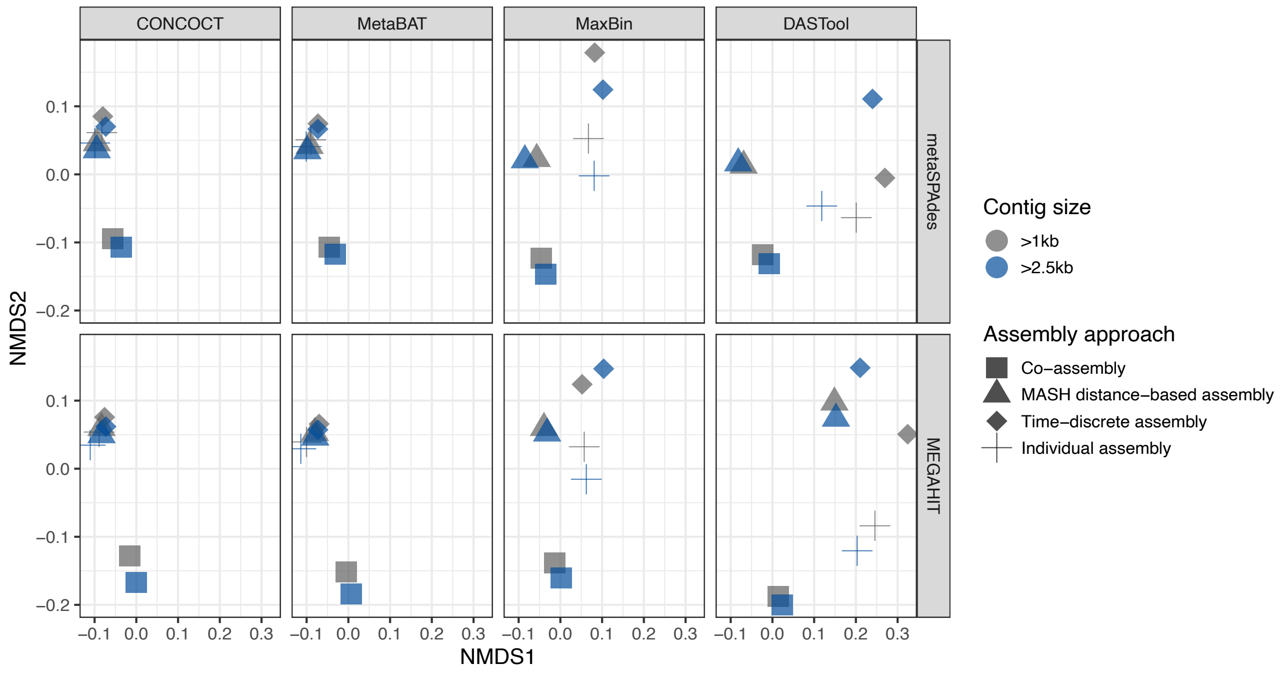


**Figure S4** Non-metric multidimensional scaling (NMDS) based on MASH distances of unrefined bins (completeness > 50%) associated with 64 assembly/binning approaches that where reconstructed with assemblies containing contigs greater than 1kbp (grey) or 2.5kbp (blue) using CONCOCT, MetaBAT, MaxBin and DAS Tool. PERMANOVA results in relation to assembly/binning approach and contig size revealed that that the assembly/binning approach explained approximately 96% (PERMANOVA, $F(31) = 46.24$, $R^2 = 0.96$, $p < .05$) of the variation, while contig size explained 2% (PERMANOVA, $F(1) = 36.16$, $R^2 = 0.02$, $p < .05$) .
