## Supplementary material for "Evaluating *de novo* assembly and binning strategies for time-series drinking water metagenomes": Figure S3

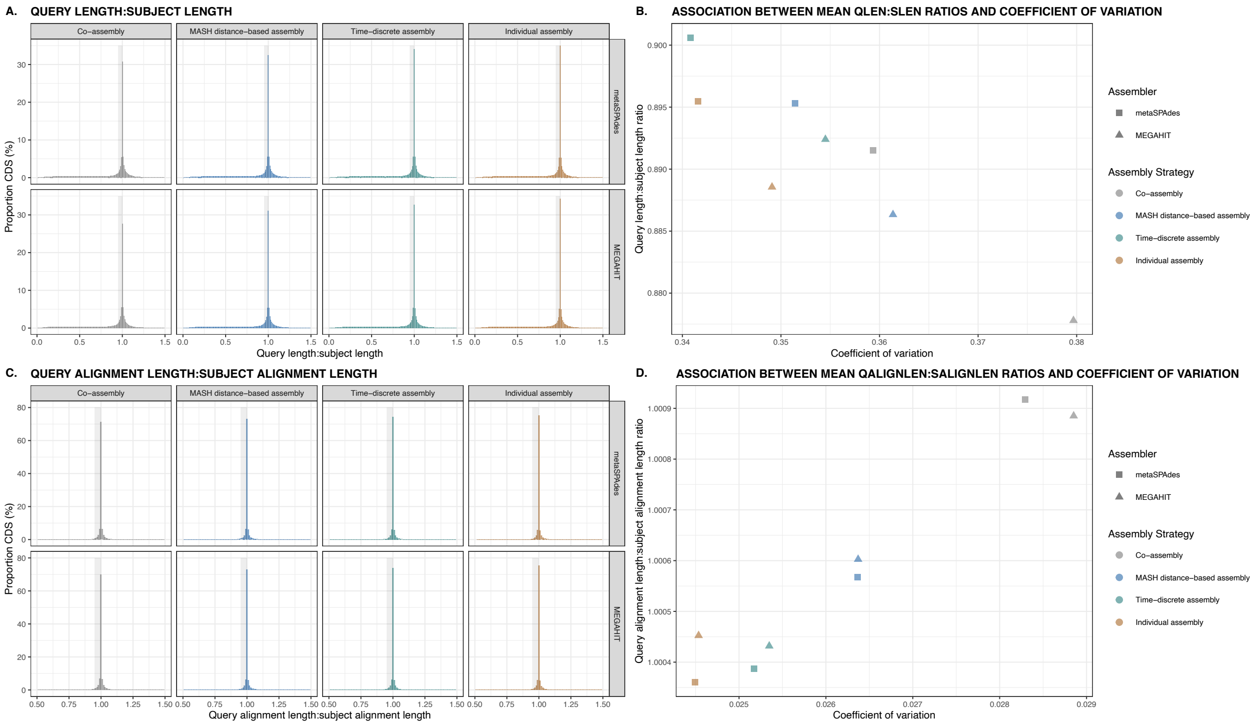


**Figure S3 A)** Histogram displaying the proportion coding DNA sequences (CDSs) of the metaSPAdes (■) and MEGAHIT (▲) assembly strategies (i.e., co-assembly of al samples (grey), MASH distance-based assembly (blue), time-discrete assembly (green), and individual assembly (orange)) with relation to their query to subject sequence length ratios (qlen/slen). The gray shade indicates qlen:slen ratios ranging between 0.95 and 1.0. **B)** Association between the mean qlen:slen ratios and coefficient of variance ($C_v$) estimates of the metaSPAdes and MEGAHIT assembly strategies. **C)** Histogram displaying the proportion CDSs of the metaSPAdes and MEGAHIT assembly strategies with relation to their query to subject sequence alignment length ratios (qalignlen/salignlen). The gray shade indicates qalignlen:salignlen ratios ranging between 0.95 and 1.0. **D)** Association between the mean qlen:slen ratios and $C_v$ estimates of the metaSPAdes and MEGAHIT assembly strategies.
