## Supplementary material for "Evaluating *de novo* assembly and binning strategies for time-series drinking water metagenomes": Figure S2

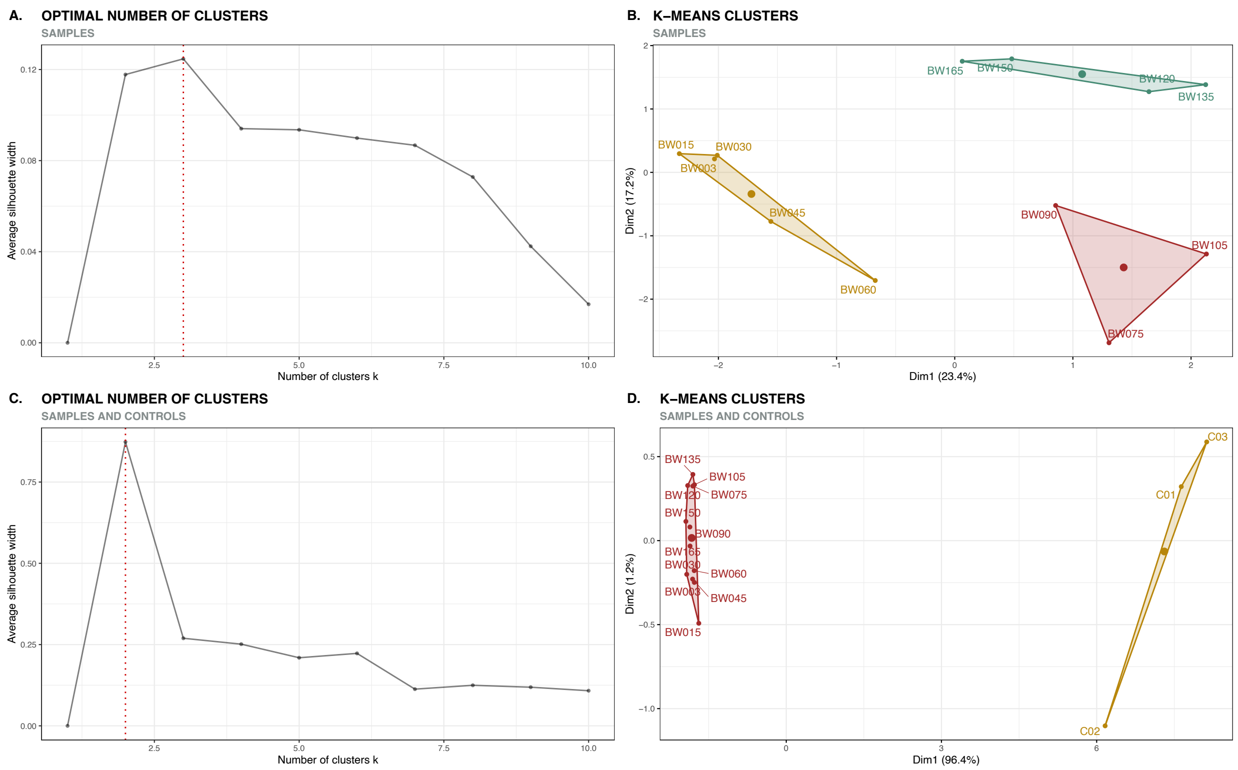


**Figure S2** K-means clustering of samples using MASH dissimilarity indices. **A)** Optimal number of clusters for samples as determined using the average silhouette method. **B)** Three distinct clusters were identified amongst samples representing BW003, BW015, BW030, BW045 and BW060 (yellow), BW075, BW090 and BW105 (red) and BW120, BW135, BW150 and BW165 (green). **C)** Optimal number of clusters for samples and controls as determined using the average silhouette method. Three distinct clusters were identified amongst samples representing BW003, BW015, BW030, BW045 and BW060 (yellow), BW075, BW090 and BW105 (red) and BW120, BW135, BW150 and BW165 (green). **D)** Two clusters grouping controls (C01, C02 and C03) independently from samples. Abbreviations: BW, bulk water; C, Controls.
