## Supplementary material for "Evaluating *de novo* assembly and binning strategies for time-series drinking water metagenomes": Figure S1

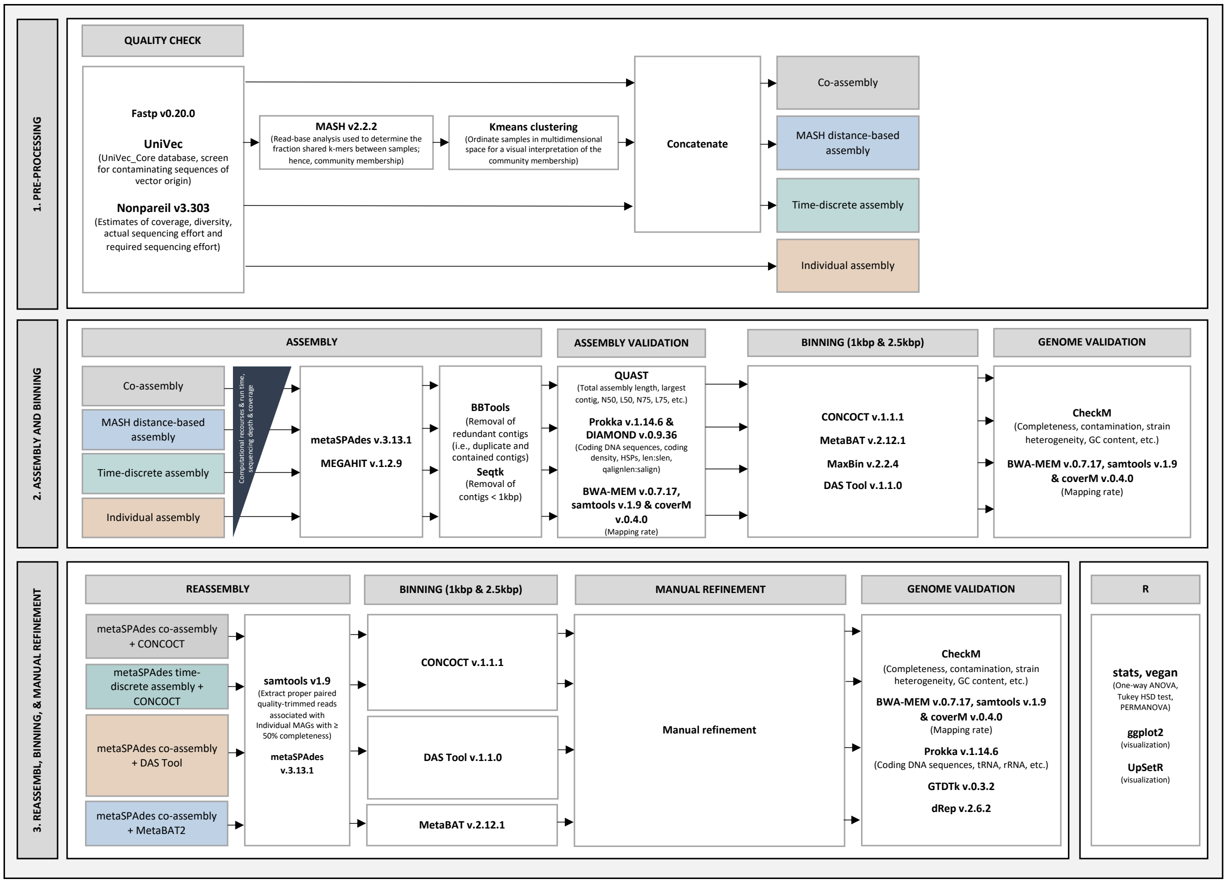


**Figure S1** Workflow used for the processing of time-series metagenomic sequencing data. In this pipeline the performance of a combination of assembly (metaSPAdes and MEGAHIT) and binning software (CONCOCT, MetaBAT2, MaxBin2, and DAS Tool) were evaluated using four assembly strategies, including individual assembly and three co-assembly approaches, i.e., co-assembly with all samples, MASH distance-based assembly, and time-discrete assembly. This resulted in 32 combinations of assembler, assembly strategy, and binning approaches (Table S2).
